# Oral and faecal viromes of New Zealand calves on pasture with an idiopathic ill-thrift syndrome

**DOI:** 10.1101/2024.12.18.629268

**Authors:** Rebecca M. Grimwood, Jessica A. Darnley, John P. O’Connell, Hayley Hunt, Harry S. Taylor, Kevin E. Lawrence, Michaela B. W. Abbott, Ruy Jauregui, Jemma L. Geoghegan

**Affiliations:** Department of Microbiology and Immunology, University of Otago, Dunedin, 9016, New Zealand; Diagnostics, Readiness and Surveillance Services, Biosecurity New Zealand, Ministry for Primary Industries, 66 Ward Street, Upper Hutt, New Zealand; Tāwharau Ora - School of Veterinary Science, Massey University, Private Bag 11222, Palmerston North 4410, New Zealand; Eltham District Veterinary Service, Eltham, 4322, New Zealand

**Keywords:** New Zealand, virus discovery, calf, idiopathic ill-thrift, enteritis, stomatitis, *Pseudocowpox virus*, *Bovine bopivirus*

## Abstract

Since 2015, an idiopathic ill-thrift syndrome featuring diarrhoea and, in some cases, gastrointestinal ulceration has been reported in weaned New Zealand dairy calves. Similar syndromes have been described in the British Isles and Australia, but investigations in New Zealand have yet to identify a specific cause. Notably, the viromes of affected calves remain understudied. We conducted metatranscriptomic analyses of oral and faecal viromes in 11 calves from a dairy farm in Taranaki, New Zealand experiencing an outbreak of this syndrome. This included nine symptomatic calves. Our analysis identified 18 bovine-associated viruses across two DNA and three RNA viral families, including five novel species. Oral viromes were dominated by *Pseudocowpox virus*, detected in all calves with oral lesions. Faecal viromes were more diverse, featuring adenoviruses, caliciviruses, astroviruses, and picornaviruses. *Bovine bopivirus*, from the *Picornaviridae* family and previously unreported in New Zealand, was significantly associated with calves showing oral lesions and diarrhoea, indicating a possible link to disease, though its role remains unclear. The diverse viral communities in the calves complicates identifying a single causative agent. Importantly, no novel viruses were significantly associated with the syndrome, and the viromes closely resembled those found in cattle globally. These findings suggest the syndrome likely has a multifactorial origin involving nutritional, management, and environmental factors rather than being driven primarily by known or novel viruses. Further research across regions and seasons is recommended to clarify the role of viruses in idiopathic ill-thrift among New Zealand calves.

## Introduction

Cattle (*Bos taurus*) play an important role in New Zealand’s economy, and were first introduced in the early 1800s with imports from Australia and the United Kingdom (1). Live cattle imports continued for many years from multiple countries, including Chile (1) and the United States (2), however, no live cattle have been imported into New Zealand since 2013 and animal product imports are subject to strict biosecurity controls. New Zealand is now the world’s largest exporter of dairy, which contributes over USD$6 billion to its economy annually (3). Dairy cows in New Zealand rely predominantly on a grass-based diet and are therefore bred on a seasonal basis to coincide dietary requirements of milk production with suitable pasture growth (2).

An idiopathic ill-thrift syndrome in weaned dairy calves of between three and 12 months of age has been reported in the United Kingdom and the Republic of Ireland (where it is termed “summer scour syndrome”) and Australia (referred to as “upper alimentary tract ulcerative syndrome” (UAUS)) between 2015 and 2019 (4–6). The ill-thrift is accompanied by diarrhea (scour) in a proportion of the calves, with a smaller proportion having severe erosive or ulcerative oral and oesophageal lesions (7). Notably, the condition is unresponsive to empirical treatments and investigations have ruled out common parasitic, bacterial, or viral agents as a cause (5). The New Zealand syndrome observed during the same period has tentatively been referred to as “calf ulcerative stomatitis (CUS)” given the presence of oral lesions in a proportion of the affected calves (8). Outbreaks of CUS in New Zealand have been extensively investigated and exotic vesicular disease (foot-and-mouth disease) has been conclusively excluded in all cases on clinical and epidemiological grounds. Further to this, samples from a number of outbreaks have been submitted to the Animal Health Laboratory (New Zealand Ministry for Primary Industries) where molecular and serological diagnostics for Foot and Mouth Disease (FMD) have all returned negative results.

Multidisciplinary investigations involving faecal examination and culture, serology, molecular assays, virus isolation, necropsies, biopsies and histological or transmission electron microscopy examination of tissues have excluded likely endemic differentials, including parasitic gastroenteritis, coccidiosis, salmonellosis, bovine viral diarrhea (BVD), calf diphtheria (*Fusobacterum necrophorum*), and malignant catarrhal fever. Investigations into the syndromes overseas have similarly excluded these differential diagnoses (6). While poxviruses have been found in a number of cases with upper gastrointestinal tract lesions, including in New Zealand, these are considered to be secondary infections in immunocompromised calves. Furthermore, poxviruses in cows are rarely linked to severe disease (9). In Australia, metatranscriptomics on oral swabs and oesopharyngeal samples from confirmed UAUS cases did not identify any novel or consistently associated viral agents. (4). Pre and post weaning calf feeding and management practices leading to poor rumen development and dysfunction have been proposed as playing a role (7). These along with environmental stressors and secondary opportunistic infections have led to the suggestion of a multifactorial aetiology (6,10).

New Zealand has maintained freedom from many pathogenic diseases of cattle, such as *Foot and mouth disease virus* (FMDV), and has successfully eradicated other pathogens including anthrax (*Bacillus anthracis*) and bovine brucellosis (*Brucella abortus*) (11). Other significant infectious diseases such as bovine tuberculosis (*Mycobacterium bovis*) are controlled through various management and eradication programmes (12). Viral diseases of cattle are generally well studied and characterised globally due to the economic importance of these animals (13–16). The emergence or introduction of infectious disease remains a concern, however, New Zealand maintains a comprehensive biosecurity and surveillance system to ensure that cattle industries in New Zealand remain free of potentially exotic or detrimental disease outbreaks (17).

We applied metatranscriptomics to investigate oral and faecal samples collected from 11 calves in a mob of 100 on a dairy support farm in Taranaki, New Zealand, experiencing an outbreak of an idiopathic ill-thrift syndrome – the seventh outbreak on the farm in eight years. Samples were collected from calves with and without oral lesions and/or diarrhea, with a view to characterising their viral communities and identifying those potentially significantly associated with the syndrome.

## Methods

### Animal ethics

Samples were collected as part of a disease investigation under the New Zealand Biosecurity Act 1993 so animal ethics approval was not required.

### Sample collection and disease characterisation

The July – August 2023 born, Friesian – Jersey cross dairy replacement heifers had been moved to the support farm from three dairy platforms from early November onwards. On 7 December, the farm manager observed that one mob of 100 calves had, as a whole, lost body condition despite being on good pasture and supplemented with concentrate feed. A greater proportion of the calves in this mob was observed to have fecal staining and or matting on the perineum and/or tail compared to another similar age mob on the farm. Over the course of one week, 14 calves from the affected mob were determined to be sick and drafted out by farm staff for closer examination and isolation in a hospital paddock. When examined, a number of these calves were noted to have oral lesions.

On 15 December an MPI incursion investigator visited the farm along with the farm veterinarian. Five healthy (control) calves as assessed by farm staff were selected for examination. Two of these calves (Y18 and Y14) had tongue lesions while a third calf (Y52) had evidence of diarrhea (tail matting). Six calves from the hospital paddock mob of 14 were also selected and presented by farm staff. All six calves had oral lesions. Calves had lesions on the tongue (n = 5), cheek (n = 3) and dental pad (n = 2). Calf W64 only had lesions on the dental pad. Five calves had tail matting or perineal staining. One calf (P18) had lesions on the external surface of the lower lip. Overall, three had oral lesions only, one had diarrhoea only, five had oral lesions and diarrhoea, and two were normal on clinical examination (Table 1).

**Table 1.** Summary of lesions in the control and case calves.

| Clinical examination | Controls | Cases | Total |
| --- | --- | --- | --- |
| Normal | 2 | 0 | 2 |
| Oral lesions only | 2 | 1 | 3 |
| Diarrhoea only | 1 | 0 | 1 |
| Oral lesions and diarrhoea | 0 | 5 | 5 |
| Total | 5 | 6 | 11 |

Oral and rectal swabs were collected from these 11 calves. In collecting swabs from calves with oral lesions, the lesions were vigorously swabbed. Swabs were placed in DNA/RNA Shield (Zymo Research), kept chilled, and placed in a -20 degrees Celsius freezer within six hours and transferred to -80 degrees Celsius freezer withing 72 hours. Under sedation and local anesthetic, lesions from calves Y17 and P18 were biopsied. Samples were fixed in 10% neutral formalin and processed routinely for histology by a commercial veterinary diagnostic laboratory.

### RNA extraction and sequencing

Frozen oral and faecal swabs stored in DNA/RNA Shield were defrosted and, using sterile forceps, transferred to BashingBead lysis tubes (0.1 and 0.5 mm) (Zymo Research) filled with 750µL of fresh DNA/RNA Shield. Samples underwent a total of five minutes of lysis in a Mini-BeadBeater-24 disruptor (Biospec Products Inc.) at 0.5 – 1.5-minute increments, with 30 seconds on ice in between. 200µL of supernatant was transferred to a sterile microfuge tube and incubated with Proteinase K. Total RNA was extracted and purified following the ZymoBIOMICS MagBead RNA kit protocol (Zymo Research), including the optional DNase I incubation step to remove DNA contaminants. RNA was quantified using a NanoDrop Spectrophotometer (ThermoFisher) and each of the 22 samples (11 oral and 11 faecal) were subject to total RNA sequencing. The Illumina Stranded Total RNA Prep with Ribo-Zero plus kit (Illumina) was used for library preparation. Libraries were sequenced on the Illumina NovaSeq 6000 platform and 150 bp paired-end reads were generated.

### Transcriptome assembly, annotation and abundance estimation

Forward and reverse reads from the 22 sequencing libraries were trimmed using Trimmomatic (v0.38) (18) to remove Nextera paired-end adapters. Bases were cut if they fell below a quality of five using a sliding window approach or if below a quality of three at the beginning or end of the reads. Library quality was assessed with FastQC (https://www.bioinformatics.babraham.ac.uk/projects/fastqc/). Trimmed reads were then assembled *de novo* using MEGAHIT (v1.2.9) (19) to generate a metatranscriptome for combined oral samples and for combined faecal samples.

Assembled contigs were annotated using a sequence similarity-based approach. Contigs were annotated at the nucleotide level against NCBI’s nucleotide (nt) database using the BLASTn algorithm (20) and at the protein level against the non-redundant (nr) protein database using DIAMOND BLASTx (v2.02.2) (21) with the “more-sensitive” flag set. Both searches were run with e-value cutoffs of 1×10^-5^. Contig abundances were estimated by mapping sequencing reads from each of the 11 individuals to the oral and faecal metatranscriptomes using Bowtie2 (22). The outputs were merged into tables containing annotated contigs and abundances for each calf. Read counts were standardised to reads per million (RPM) by first dividing contigs counts in each library by their respective library size and multiplying these by one million.

### Bovine virus discovery

Bovine-associated viruses were screened for manually by filtering contigs by those annotated as being from the Viruses domain/superkingdom. Viral sequences with e-values <1×10^-10^ were inspected with additional BLASTn and BLASTp searches using the online BLAST tool to eliminate false positive hits and potentially endogenous viral elements (EVEs). Putative viruses were considered if their top BLASTp genetic matches were to previously known vertebrate host-infecting viruses or those known to infect cattle (*Bos*).

### Phylogenetic analysis of bovine viruses

RNA-dependent RNA polymerases (RdRps), DNA polymerases, or hexon proteins (in the case of adenoviruses where no polymerases were identified) from putative viruses identified in the calf metatranscriptomes were collated and organised by their proposed viral families for phylogenetic analysis. Viruses were placed into an evolutionary context using maximum-likelihood phylogenetic trees generated using IQ-TREE (v1.6.12) (23). First, representative viral sequences from each genus in the viral families identified were collected from NCBI’s Taxonomy database (https://www.ncbi.nlm.nih.gov/taxonomy) and amino acid translations of these were aligned with the bovine-associated viruses identified here using the L-INS-i algorithm in MAFFT (v7.450) (24). Alignments were manually inspected to observe the alignment of conserved motifs and ambiguously aligned regions were trimmed using trimAL (v1.2) (25) with the “automated1” flag set. Then, IQ-TREE was used to generate phylogenies based on these alignment with the ModelFinder “MFP” flag set to select the best-fit model of amino acid substitution and 1000 ultra-fast bootstrapping replicates (26). The “alrt” flag was also added to perform 1000 bootstrap replicates for the SH-like approximate likelihood ratio test (27). The resulting phylogenetic tree was rooted at its midpoint and annotated in FigTree (v1.4.4) (http://tree.bio.ed.ac.uk/software/figtree/). Viruses were determined to be novel if they shared less than 90% amino acid identity with other known viruses.

### Differential viral abundance analysis

To determine which viral groups were more abundant in calves with oral lesions compared to calves without oral lesions, a differential abundance analysis was performed using the DESeq2 package in R (v4.11) with eight “affected” calves and three “unaffected” calves as controls (Table 1, Figure 2B). We also repeated the analysis between calves with (five) and without (six) diarrhoea and those with both key symptoms (oral lesions and diarrhoea) (four) versus those with none or only one symptom (seven). Total virome abundances (as raw reads) of the calves were condensed at genus level where possible, followed by family, order, or as “Unclassified Riboviria” (RNA viruses) or “Unclassified DNA viruses” where more detailed taxonomic information was not available. Viral groups were considered significantly more abundant in affected calves when the FDR-adjusted p-values (q-values) were < 0.05. Viral abundance differences were considered in terms of their change in unaffected calves. The Benjamini-Hochberg (BH) multiple correction procedure was used to re-adjust p-values.

The initial categorisation of the calves into healthy and unhealthy groups, when sampled by MPI in December 2023, was based on a diagnosis made by farm staff. However, the farm staff were untrained and were unlikely to have carried out thorough oral examinations and were focused on removing the most severely affected calves to the hospital paddock. As such, we feel that recategorising the calves into affected and unaffected, based on the results of the clinical examination carried out by the veterinarians (Supplementary Table 1), was justified to broadly compare the viromes of more severely affected calves to those without oral lesion.

## Data availability

Sequencing reads are available on the NCBI Sequence Read Archive (SRA) under the BioProject accession (PRJNA1164330) and virus sequences are available under the GenBank accessions (PQ583780 – PQ583802). R scripts and additional viral sequence data can be found on GitHub: https://github.com/maybec49/NZ_Calf_Viromes.

## Results

### Histology of oral lesions

Histologically, oral lesions were characterised by multifocal areas of ulceration (Figure 1), covered by crusts composed of neutrophils, karyorrhectic debris and fibrin (Figure 1C and 1E). There was hyperplasia (thickening) of the adjacent intact mucosa (Figure 1C), with variable degrees of ballooning degeneration of keratinocytes in the stratum corneum and stratum spinosum (Figure 1D). Occasional rounded, hypereosinophilic keratinocytes were present in the intact mucosa, consistent with apoptotic cells (Figure 1D). There were no apparent viral inclusion bodies in the sections examined. These findings are consistent with previous cases of idiopathic ill-thrift in calves in New Zealand (H. Hunt, pers. comm.)

**Figure 1.**
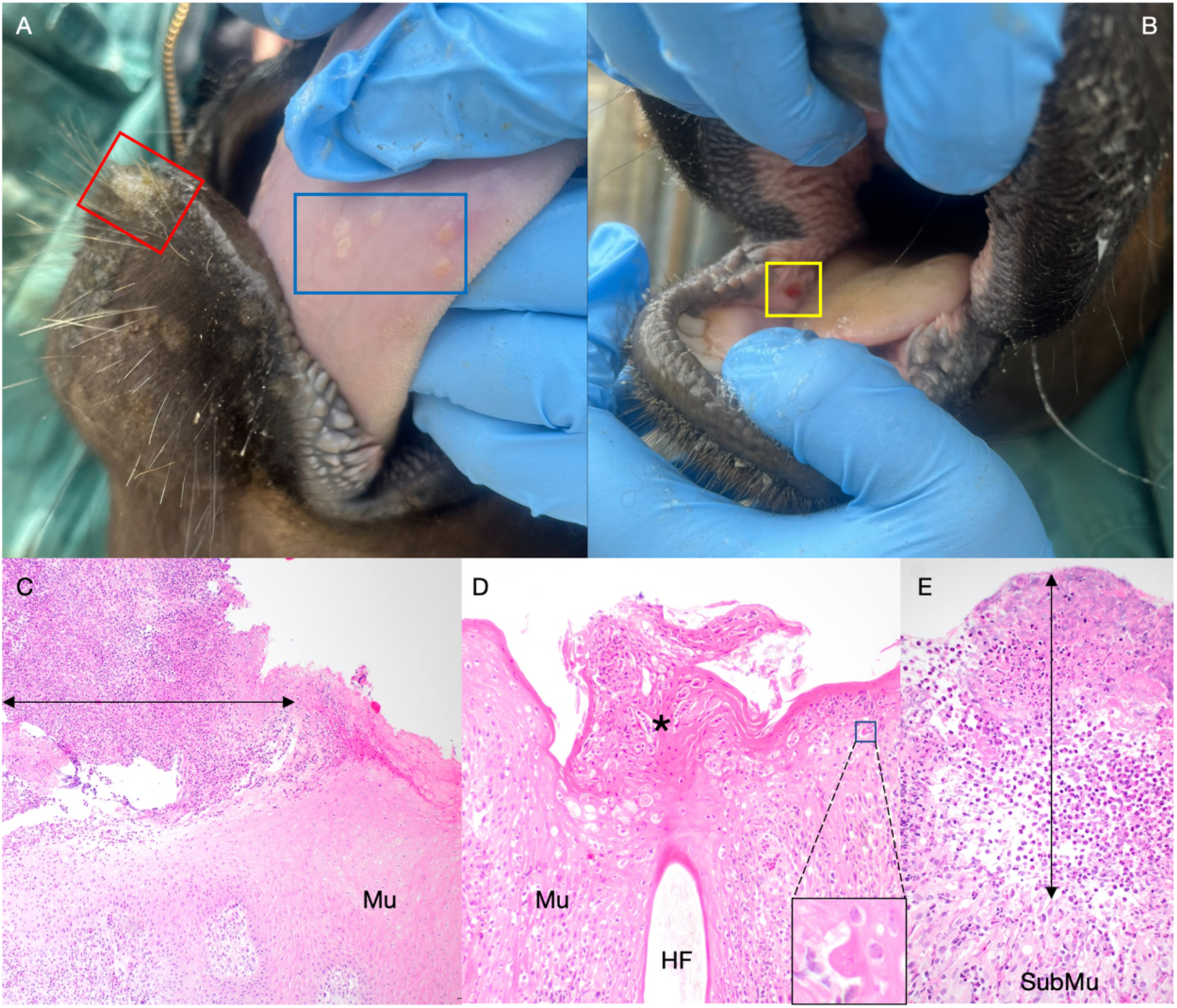
Clinical and histological appearance of oral lesions in calves with idiopathic ill-thrift. **A.** Multifocal pale circular lesions on the underside of the tongue (blue box) and an area of thickened skin on the lower lip (red box). **B.** A focal area of ulceration in the mucosa of the right lower cheek. **C.** Histology of the tongue lesions in the blue box in image A. There is an area of ulceration (arrow) which is covered by a thick crust composed of degenerate neutrophils, fibrin and karyorrhectic debris. The adjacent mucosa (Mu) is hyperplastic. **D.** Histology of the lip lesion in the red box in image A. The mucosa (Mu) is hyperplastic and there is marked ballooning degeneration of keratinocytes in the stratum corneum and spinosum. Overlying a hair follicle (HF), there is hyperkeratosis and sloughing keratocytes are admixed with small numbers of neutrophils (*). There are occasional hypereosinophilic, rounded keratinocytes (apoptotic cells), as shown in the inset. **E.** Histology of the cheek lesion in the yellow box in image B. There is no identifiable mucosa as it is ulcerated and there are large numbers of neutrophils mixed with fibrin in its place (arrow). Inflammatory cells extend into the submucosa (SubMu) and are accompanied by loose connective tissue proliferation (granulation tissue).

### Sequencing and total viromes

Metatranscriptomes for each of the two sample types were assembled from all 11 calves, producing 381,570 oral-associated contigs and 2,834,178 faecal-associated contigs. Viruses accounted for an average of 10% of oral reads and 45% of the faecal reads across the calves (Table 2). Bovine-associated viral reads across the two sample types accounted for 113 to 8,848 RPM.

**Table 2.** Calf sequencing library and metatranscriptome details and virome abundances.

| Calf ID | Oral lesions | Oral reads | Faecal reads | Total oral virome | Total faecal virome | Total bovine viruses RPM |
| --- | --- | --- | --- | --- | --- | --- |
| P18 | Y | 18,381,405 | 14,705,921 | 9% | 53% | 1072 |
| W64 | Y | 12,838,378 | 13,216,843 | 2% | 32% | 113 |
| Y14 | Y | 15,485,106 | 12,354,419 | 14% | 35% | 447 |
| Y17 | Y | 14,220,387 | 11,868,221 | 8% | 91% | 8849 |
| Y18 | Y | 16,365,398 | 13,717,264 | 14% | 29% | 1466 |
| Y35 | Y | 13,484,657 | 14,014,061 | 9% | 36% | 435 |
| Y41 | Y | 18,880,150 | 11,767,740 | 7% | 25% | 1660 |
| Y70 | Y | 32,056,352 | 12,210,744 | 9% | 38% | 1180 |
| W3 | N | 20,375,283 | 10,345,261 | 4% | 51% | 204 |
| Y7 | N | 15,275,728 | 11,823,969 | 23% | 62% | 180 |
| Y52 | N | 13,268,730 | 13,219,497 | 2% | 46% | 97 |

We identified viruses from two DNA viral families (*Adenoviridae* and *Poxviridae*) and three RNA viral families (*Astroviridae*, *Caliciviridae* and *Picornaviridae*) in the calve samples (Figure 2C). *Picornaviridae* was the most widespread family, with 91% of oral samples and 100% of faecal samples containing picornaviral sequences. Astroviruses were also present in all faecal samples.

**Figure 2.**
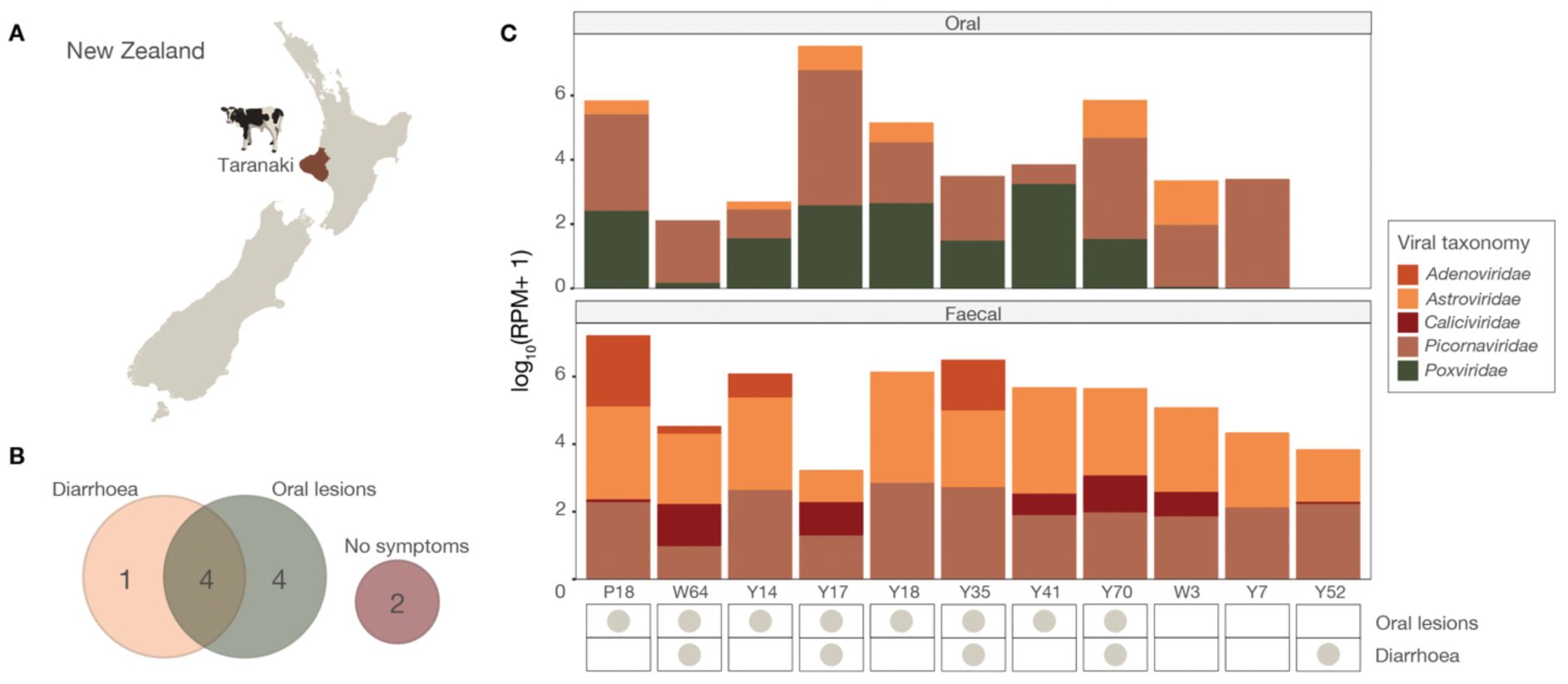
Summary of calves with CUS symptoms and abundance of viral families. A) Map of New Zealand with Taranaki highlighted. B) Venn diagram of the number of calves affected by ill-thrift symptoms. C) Bar plot of log_10_-transformed RPM counts of viral families in the 11 calves. Viral families present in oral samples are shown in the top panel and those in faecal samples are shown on the bottom panel. Chart showing which symptoms the calves were affected by is shown below.

*Poxviridae* was the secondmost abundant family in oral samples and was only identified in calves with oral lesions. Adenoviruses and caliciviruses were also primarily found in faecal samples from affected calves, but were only associated with 50% and 63% of affected calves, respectively.

Six genera within the *Picornaviridae* were identified across the samples (*Apthovirus* (bovine rhinitis viruses), *Bopivirus*, *Enterovirus*, *Hunnivirus*, *Kobuvirus*, and *Parechovirus*). Of these, enteroviruses, kobuviruses, and bovine rhinitis viruses, as well as mammalian astroviruses (mamastroviruses) were the most widespread and abundant across the samples, being found in both affected and unaffected calves (Figure 3 and Supplementary Figure 1). Pseudocowpox virus (genus: *Parapoxvirus*), known to be infecting calves from Taranaki, was also identified only in calves affected by oral lesions at relatively high abundance (1.4 – 465 RPM in oral samples). The genera *Nebovirus* (family: *Calciviridae*), *Mastadenovirus*, *Bopivirus*, *Hunnivirus*, and *Parechovirus* were either absent or present at very low abundances in unaffected calves. On average, calves harboured 7.2 viral genera (affected: 7.5 and unaffected: 6.3) (Figure 3 and Supplementary Figure 1).

**Figure 3.**
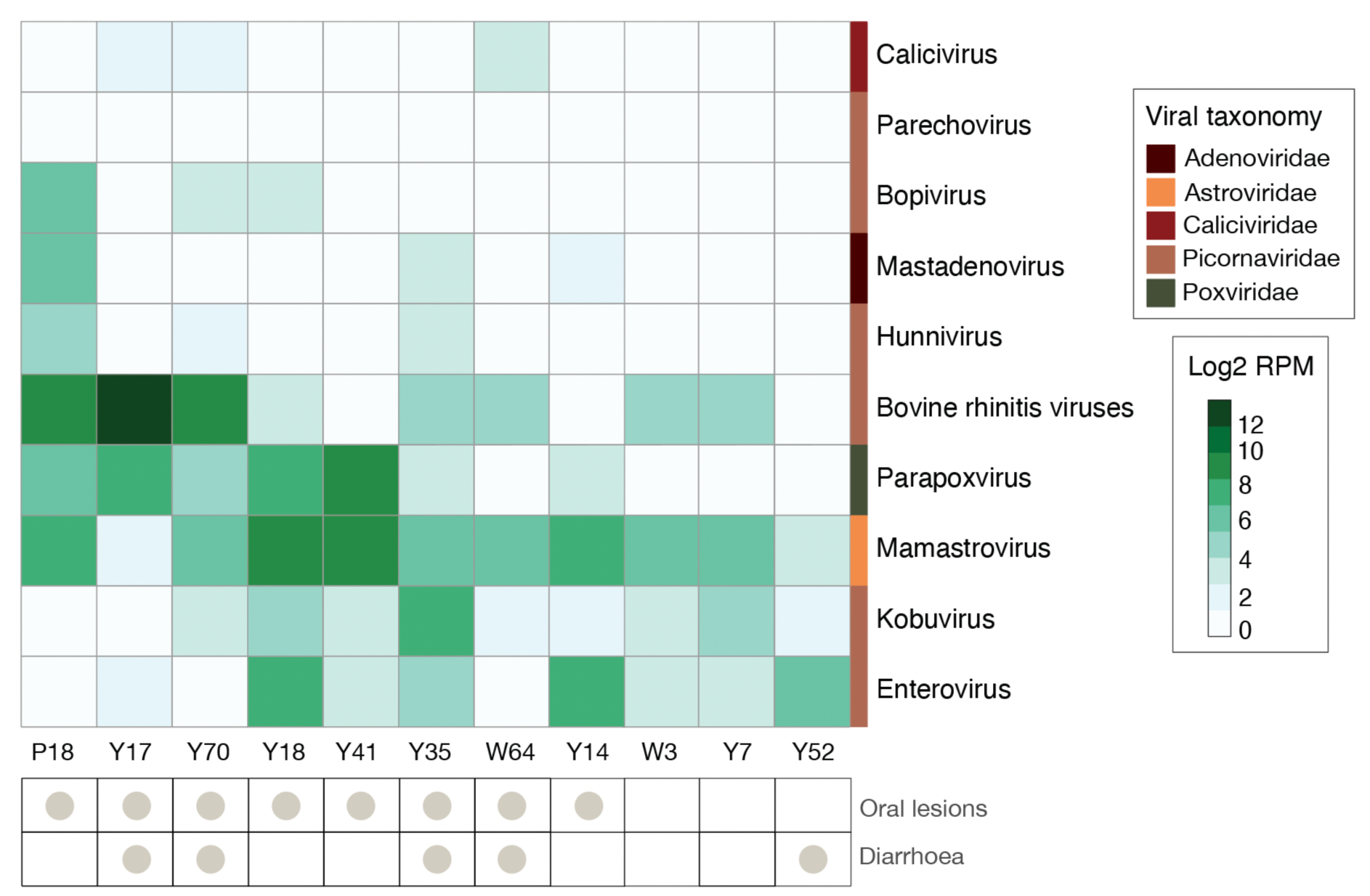
Heatmap of bovine-associated viral genera and species per sample. Abundances are shown as log2-transformed RPM with white indicating zero abundance and darker green indicating higher abundance. Chart showing which symptoms the calves were affected by is shown below the heatmap.

### Phylogenetic analysis of bovine viruses

To determine the evolutionary histories and novelty of the bovine-associated viruses identified in the calves, we estimated phylogenies using polymerase or hexon proteins from the identified species (Figure 4 and Table 3). Five viral families, represented by 18 viruses, could be assessed: the *Adenoviridae* (n=1), *Poxvirida*e (n=1), *Astroviridae* (n=2), *Caliciviridae* (n=1), and *Picornaviridae* (n=13). Of these species, five were presumed to be novel and are described in the following sections.

**Figure 4.**
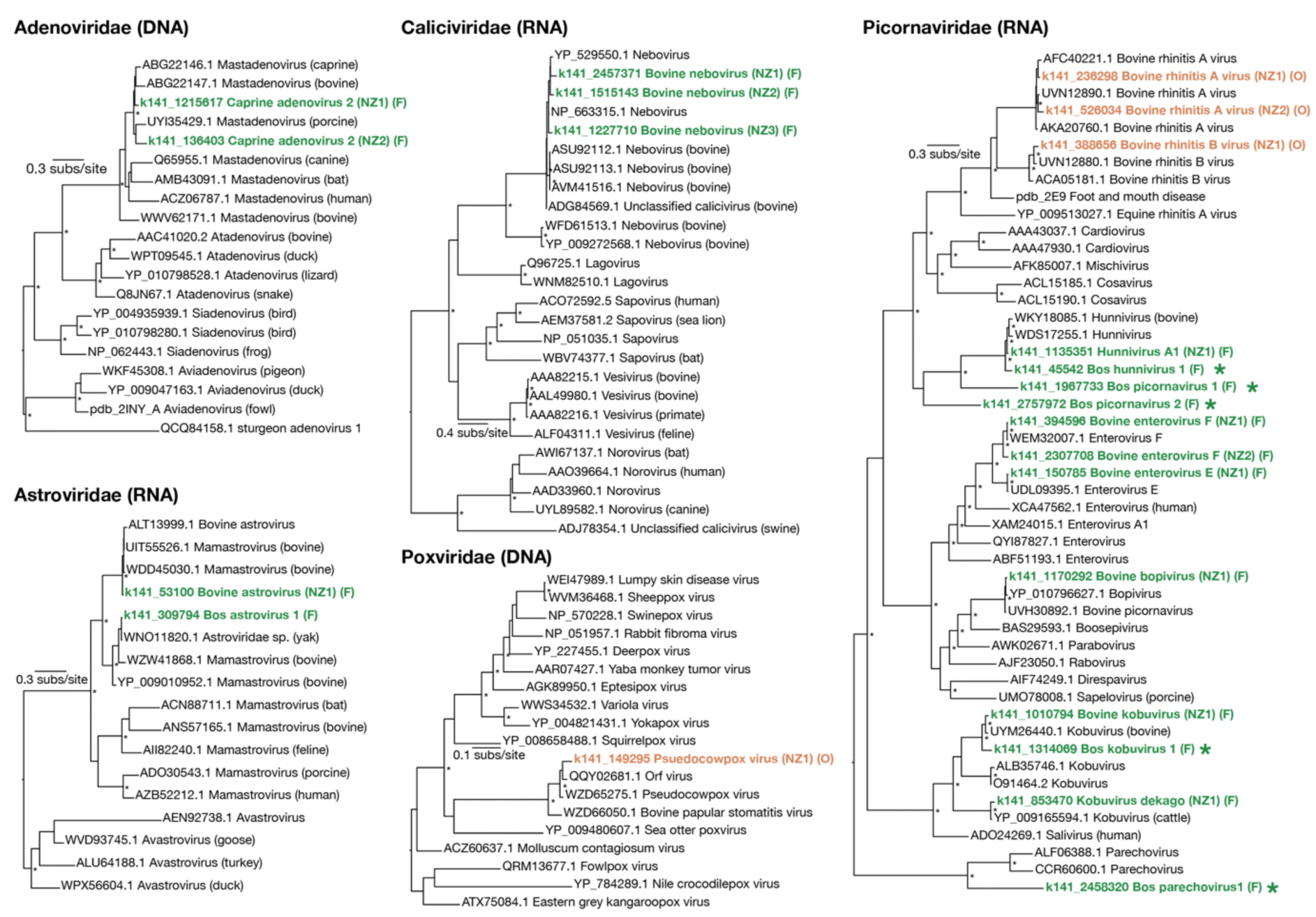
Maximum likelihood phylogenetic trees of bovine-associated RNA and DNA viruses present in New Zealand calves based on DNA and RNA polymerase sequences. Viruses identified in oral samples (O) are indicated in orange and viruses identified in faecal samples (F) are highlighted in green. Viral polymerases sharing less than 90% amino acid identity with their closest relative are indicated by large asterisks. Trees are rooted at their midpoints and node UF-bootstrap values ≥ 95% are indicated by small asterisks at the node.

**Table 3.**
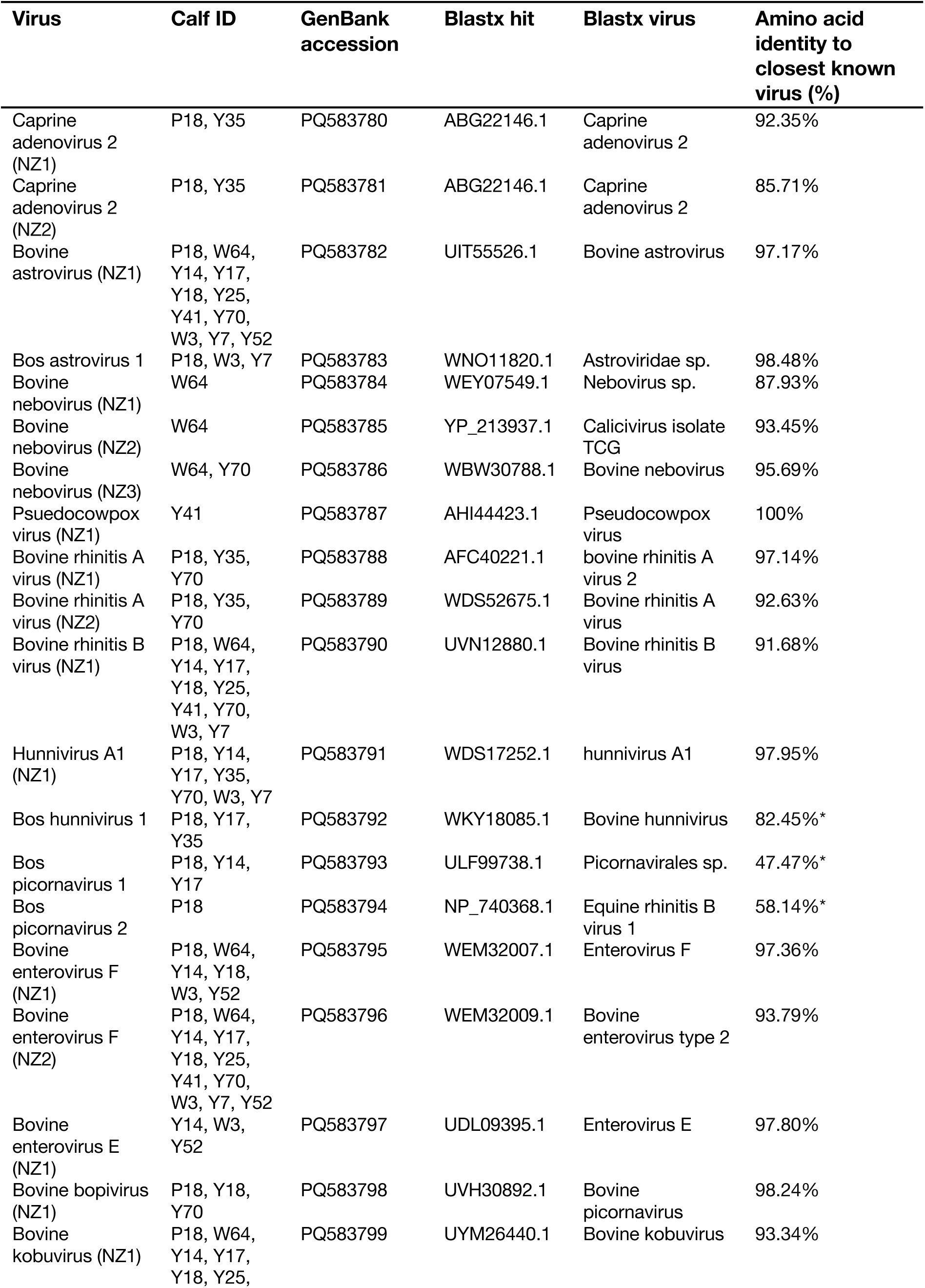
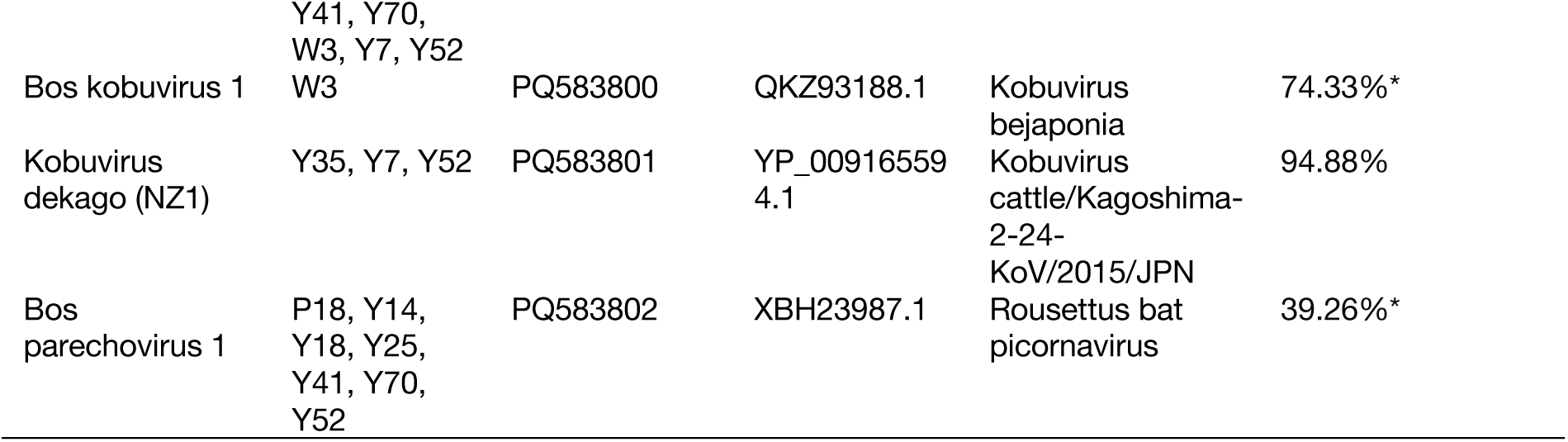
Summary of bovine viral polymerases and closet viral relatives identified in the calves.

### Oral viruses

Three viral species were identified in oral samples – *Pseudocowpox virus* (family: *Poxviridae*) and bovine rhinitis viruses A and B (family: *Picornaviridae*; genus: *Apthovirus*). The RPO147 subunit of the poxvirus identified in these samples shared 100% amino acid identity with *Pseudocowpox virus* (AHI44423.1) from cattle exhibiting oral lesions in Bangladesh (28), indicating the virus from the Taranaki calves is the same species. Bovine rhinitis viruses are common viruses of cattle causing respiratory symptoms (29), including being established causes of bovine respiratory disease (BRD) (30). We identified viral sequences sharing at least 91% amino acid sequence identity with both *Bovine rhinitis virus A* and *Bovine rhinitis B* from the *Aphthovirus* genus. The apthovirus polymerases from the calves shared 92-97% identity with these rhinitis viral species (Table 3), suggesting they are the same viral species. *Bovine rhinitis virus B* was identified in 91% (10/11) of the oral samples, while *Bovine rhinitis virus A* was only found in three.

### Faecal viruses

We identified two adenovirus hexon proteins across the faecal metatranscriptome, both phylogenetically clustering with mastadenoviruses (mammalian adenoviruses) from bovine, caprine, and porcine hosts (Figure 4). The hexons shared 86-92% amino acid sequence identity with *Caprine adenovirus 2* (ABG22146.1). These viruses are frequently found in the intestinal tracts of ruminants (31) and some adenoviruses can cause diarrheal or respiratory illness in calves (32). Since these hexons were closely related and found across the same two calves, they likely represent different segments of the same viral species. Astroviruses were identified in all 11 faecal samples. Two astroviral RdRps fell into distinct clades and shared 97 to 98% amino acid identity with astroviral proteins identified in yaks (WNO11820.1) and calves with diarrhea in China (*Bovine astrovirus*) (33), therefore representing previously known astroviruses. We propose the more descriptive name of Bos astrovirus 1 for the undefined *Astroviridae* sp. from yak (WNO11820.1) here. Three RdRp segments from neboviruses (family: *Caliciviridae*) were found in a small number of the faecal samples. These shared 88 to 96% amino acid identity with other bovine neboviruses, which have been linked to diarrhea in calves (34). As all three sequences were closely related and found across the same individuals, they also likely represent different segments from a single, previously known viral species – *Bovine nebovirus*.

Finally, we identified an additional 11 picornaviruses across the faecal metatranscriptomes falling within or close to the genera *Hunnivirus*, *Enterovirus*, *Bopivirus*, *Kobuvirus*, and *Parechovirus*. We noted two hunnivirus species – one previously known, *hunnivirus A1*, originally identified in cow faecal samples from China (98% amino acid identity) and another slightly more divergent species we named Bos hunnivirus 1. Hunniviruses have been found in a range of mammalian species, including healthy and diarrheal cows (35). Kobuviruses and *Enterovirus E* and *F* were detected in between one and 11 calves and are both common intestinal viruses with global distribution (36). We also identified a previously known bopivirus, *Bovine picornavirus* (UVH30892.1). Viruses from the *Bopivirus* genus have been found in faecal samples from ruminants in Hungary, Italy, China, and the USA (37). Of the faecal viruses, five were assumed to be novel viruses sharing less than 90% amino acid identity with known viruses (range: 39 to 82%). These included the aforementioned novel hunnivirus, two bovine (*Bos*) picornaviruses falling basal to the *Hunnivirus* genus, a kobuvirus (Bos kobuvirus 1), and a bovine parechovirus, which was related to a virus isolated from faecal samples of cows from Japan (38).

### Differentially abundant viral groups in calves with oral lesions

To determine which viral groups may be more abundant and significantly associated with more severely affected calves (those with oral lesions, with or without diarrohea), we performed a differential abundance analysis on oral and faecal viromes, including all viruses identified even if they were likely to be from dietary or environmental sources (Figure 5). A total of 148 viral taxonomic groups were present in the oral viromes (Figure 5A) and 234 in the faecal viromes (Figure 5C). Of these, 25 viral groups were found to be significantly more abundant in oral samples from affected calves (Figure 5B). Only the *Parapoxvirus* genus is associated with infections in mammalian/bovine species (q-value = 1.2×10^-8^). In faecal viromes, on the other hand, five viral groups were found to be significantly more abundant in affected calves and another five more abundant in unaffected calves (Figure 5D). Again, only one viral genus that was significantly more abundant in affected calves, *Bopivirus*, is known to infect mammals (cows) (q-value = 0.03). Other viral groups significantly different between the two groups of calves included bacteriophages such as those from the *Fiersviridae* and plant-infecting virus genera including *Gammacarmovirus* and *Tombusvirus*, likely associated with calves through dietary or environmental sources. Importantly, no bovine- or vertebrate-associated viruses were found to be significantly more abundant in the oral and faecal viromes of calves affected by diarrohea compared to those without, or in those of calves with multiple symptoms as opposed to only one or none (Supplementary Data 2).

**Figure 5.**
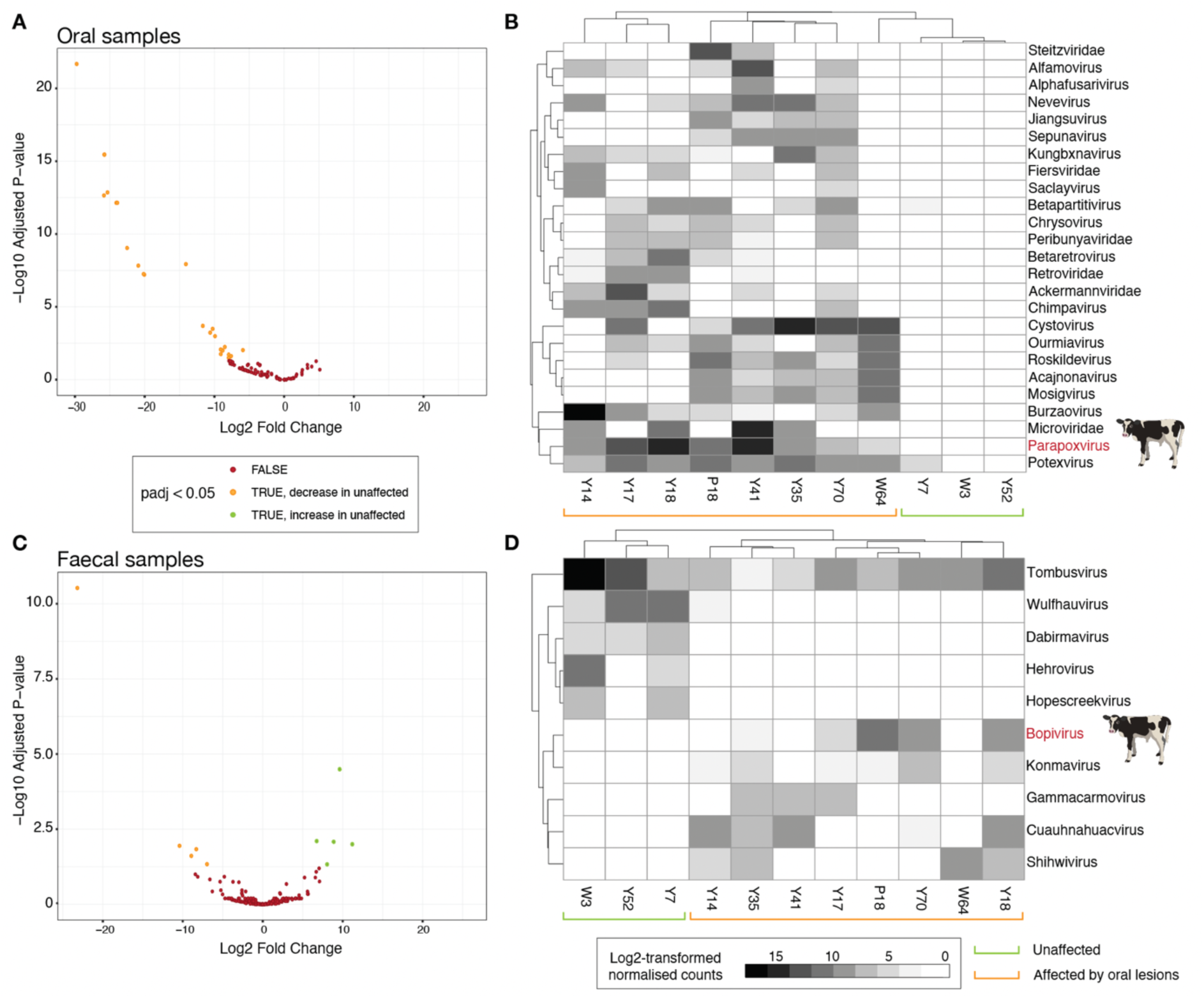
Significantly differentially abundance viral groups in the full viromes of calf oral and faecal samples were identified using DESeq2 between calves with and without oral lesions. A and C) Volcano plots showing significantly differentially abundant viruses (padj < 0.05) in oral (A) and faecal (C) samples (left). Viruses significantly more abundant in unaffected samples are represented by green dots and those significantly more abundant in affected calves are orange dots. B and D) Clustered heatmaps showing the significantly differentially abundant viral groups as log2-transformed normalised viral counts in oral swab (B) and faecal swab (D) samples (right). White squares indicate no virus detected in a sample, darker grey indicate increased viral abundance. Bovine-associated viral genera are highlighted in red and shown by a cow symbol.

## Discussion

In this investigation, we characterised the viral communities of poorly performing weaned dairy calves with clinical disease from a dairy farm in the Taranaki region of New Zealand, as well as calves without symptoms, to identify any potential viral aetiology. Overall, oral and faecal viromes consisted of a combined total of 18 bovine-associated viruses that could be placed at the species level across five DNA and RNA viral families. Coinfections were common, with all calves carrying multiple viral species. In general, the viromes of calves closely mirrored those of calves overseas (39) and did not contain any notifiable exotic pathogens. Oral viromes contained *pseudocowpox virus* and *bovine rhinitis virus* genotypes A and B. *Pseudocowpox virus* is usually associated with lesions on the udders of cows but is one of a few poxviruses that can cause oral lesions in cattle and has zoonotic potential (40). The virus is widespread and and has previously been identified on this farm. We found *pseudocowpox virus* to be significantly associated with affected calves, being found in all eight calves with oral lesions at relatively high abundance (up to 465 RPM) and none of the unaffected calves.

Reports of viral involvement in lesions seen overseas, however, have been inconsistent and the observed lesions are generally more severe than would be expected for typical poxvirus outbreaks (6). In addition, the main microscopic features of the oral lesions in the current outbreak included erosion and ulceration, whereas poxviral lesions are characterised by epithelial proliferation and hyperkeratosis (41). Bovine rhinitis viruses are common causes of respiratory disease in cattle worldwide (29,42) and have also been found to be carried asymptomatically (16). These picornaviruses are part of the same genus, *Aphthovirus*, as FMDV (29), however, they have not been associated with oral lesions and were identified in both affected and unaffected calves.

Diarrhoea is one of the most common causes of morbidity and mortality in calves, with viruses frequently being involved (36). Faecal viromes of these Taranaki calves were more diverse than oral viromes, containing adenoviruses, caliciviruses, astroviruses, and picornaviruses. Astroviruses and picornaviruses, including enteroviruses, hunniviruses, and kobuviruses, are extremely prevalent in bovine faecal samples and are frequently associated with, or causative of, diarrhea (15,36,43).

Coinfections are common and can be involved in more severe disease. For example, bovine astrovirus infections and coinfections are common but typically do not result in clinical disease on their own, rather when coinfected with other enteritically-transmitted viruses can lead to severe diarrhea and more extensive astroviral infections (39,44). Coinfections were also prevalent in the Taranaki farm, with all 11 calf faecal viromes containing at least four different viruses. Overall, however, no bovine-specific viruses were significantly more abundant in calves with diarrhoea than those without, regardless of oral lesions status. This suggests total viral burdens could potentially be contributing to disease and increased severity rather than being the result of a single agent.

Of note, we found that bovine picornavirus (bopivirus), in the genus *Bopivirus*, was also significantly associated with calves affected by oral lesions with or without diarrhoea. Bopiviruses, and related ovipi- and gopiviruses from ovine (sheep) and caprine (goats), respectively, are a relatively new set of enteric viruses found in domestic livestock and wild deer (45,46). Bovine bopivirus has not been previously described in New Zealand, making this the first report of this virus here. It is entirely plausible that bovine bopiviruses have been present in New Zealand for some time, and have been found during this present study due to the more comprehensive genomic methods used. Bovine bopiviruses have only recently been described worldwide, being found in Asia, Australia, Europe and the US to date (37,45–47). Furthermore, their role in disease pathogenesis is yet to be elucidated (45), making it difficult to interpret the clinical significance of their presence in these calves. While it is difficult to disentangle primary or multifactorial causes of diarrhea in the presence of coinfections, bopiviruses have been posited as a contributing factor to other cases of acute gastroenteritis (47). Although no single enteric virus was found consistently across all affected calves, it is possible that the bopivirus found here could play an important role in this disease.

Importantly, no notifiable exotic viral agents were identified. The four novel picornaviruses we detected were also not associated with the syndrome and were closely related to other common bovine-associated faecal viruses and may represent RNA viruses that have undergone divergence since *Bos taurus*’ introduction to New Zealand. A high diversity of RNA viruses in bovine samples is not unusual (48). Virome investigation of affected calves in Australia similarly ruled out a novel viral aetiology (4).

Seasonal differences in pathogen diversity and abundance have been observed (49–51) and these could be explored in New Zealand dairy calves in association with such events, considering breeding occurs seasonally. A limitation of this study was that all samples came from a single dairy support property in December (early Southern summer) and hence, metagenomic exploration should also be expanded to additional months and farms around New Zealand to validate the viral findings made here and help elucidate potential contributing factors to the disease.

Calf viromes in New Zealand are largely consistent with those identified overseas with little divergence and primarily harbour common faecal- and oral-associated viral species. We determined *Pseudocowpox virus* to be associated with the oral lesions seen in affected calves from Taranaki. The poxvirus is known to be both a primary pathogen and a secondary invader of damaged tissue, and although this is an important finding, further work is required to better understand the pathogenesis of oral lesions in cases of idiopathic ill-thrift in calves. In addition, the enterically-associated bopivirus was significantly associated with affected calves and has been linked to diarrhea in other populations and species. However, calf faecal viromes contained several viral species linked to gastroenteritis that have not been posed as direct causative agents and therefore the contribution of any one or combination of these viruses to their condition should not be overstated.

## Acknowledgements.

RMG is funded by a University of Otago Doctoral Scholarship. JLG is funded by a New Zealand Royal Society Rutherford Discovery Fellowship (RDF-20-UOO-007) and the Webster Family Chair in Viral Pathogenesis. The authors would like to thank the farm owners and staff for their engagement and assisance in this investigation.

## Supplementary material

**Supplementary Figure 1.**
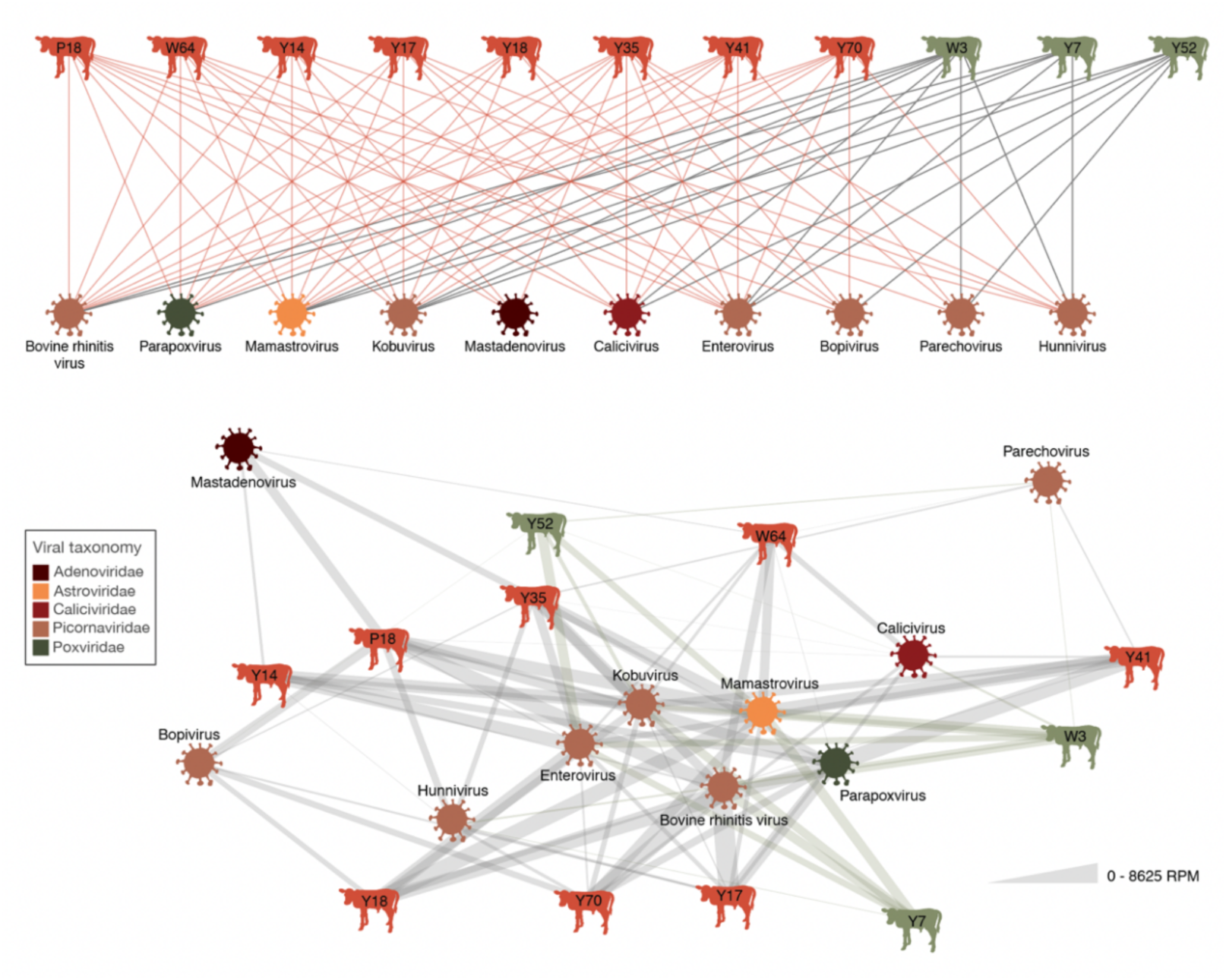
Bipartite networks showing calves and their co-infecting viral genera or species. Railway network showing individual calves and their co-occurring viral groups (top). Weighted network showing co-occurring viruses and their combined abundances in oral and faecal samples in RPM (bottom). Calves unaffected by oral lesions are highlighted in green and calves affected by oral lesions are highlighted in red.

**Supplementary Table 1.** Clinical presentations summary of the sampled dairy calves.

| Calf VID | Oral lesions? | Faecal matting or staining? | Notes |
| --- | --- | --- | --- |
| Y7 | N | N |  |
| W3 | N | N |  |
| Y18 | Y | N | “Healthy” but lesion on tongue with diphtheritic |
| Y52 | N | Y | “Healthy” but tail matted |
| Y14 | Y | N | “Healthy” but lesions under tongue |
| W64 | Y | Y | Dental pad lesion? |
| Y41 | Y | N | Clean tail. Lesion under tongue |
| Y17 | Y | Y | Lesion on lower right cheek |
| Y35 | Y | Y | Lesion on top gum |
| Y70 | Y | Y | Lesion on side of cheek, underside of tongue, top of tongue. Diphtheritic material on tongue. |
| P18 | Y | N | Pustules (?) underside of tongue. Ulcers on frenulum. Bovine respiratory syncytial virus? |

