## Supplementary figures and images for "Oral and faecal viromes of New Zealand calves on pasture with an idiopathic ill-thrift syndrome"

### Supplementary Figure 1

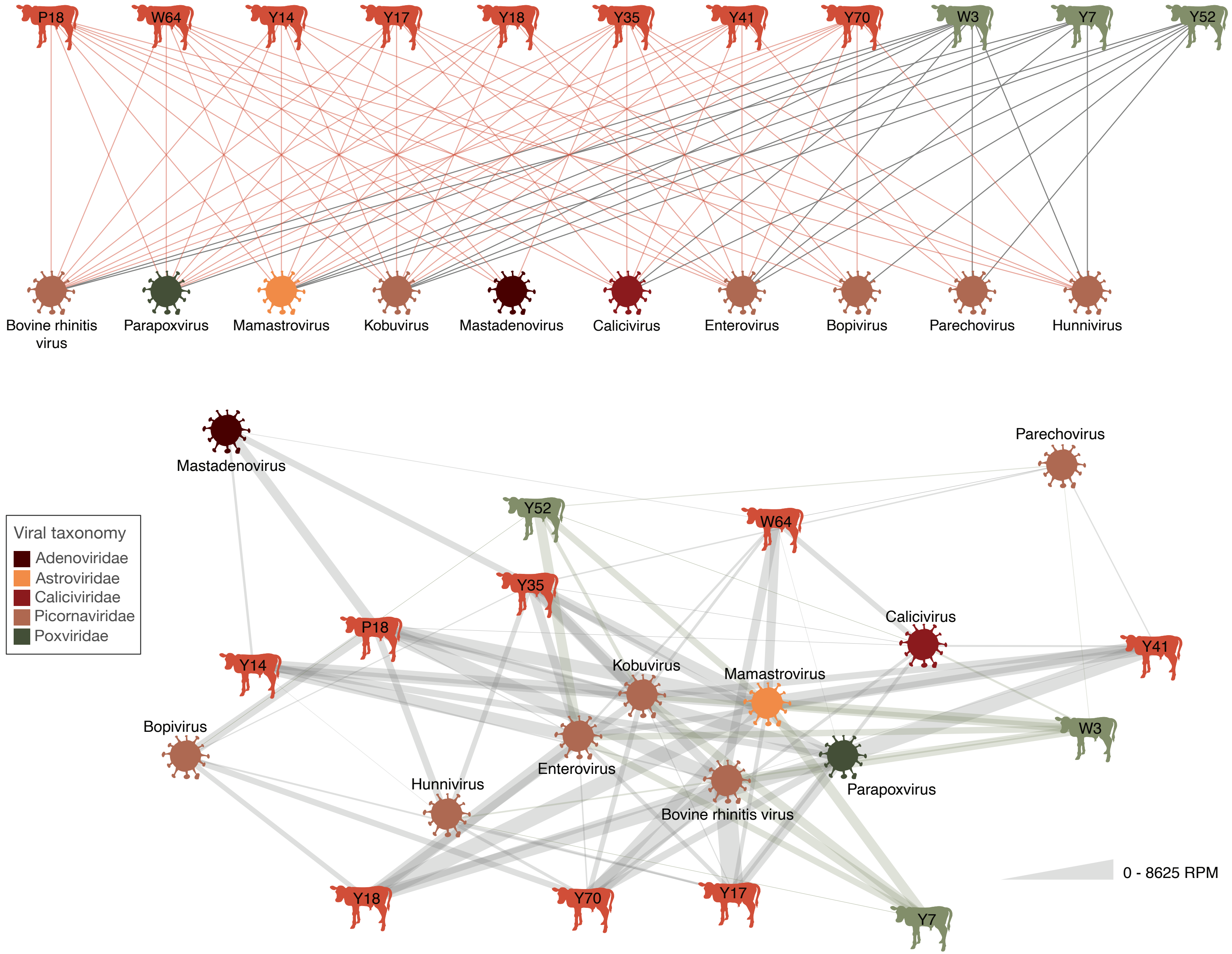
